# Influence of fatigue on reaction times and corticospinal excitability during movement preparation

**DOI:** 10.1101/2020.05.20.105742

**Authors:** William De Doncker, Katlyn E Brown, Annapoorna Kuppuswamy

## Abstract

Slower self-selected ballistic movement speeds and reduced corticospinal excitability at rest are associated with post-stroke fatigue (PSF). It is unclear if fatigue-related differences in corticospinal excitability, measured using transcranial magnetic stimulation, prior to a movement will explain slower movement speeds. We hypothesized that the levels of PSF explains the modulation of corticospinal excitability during movement preparation and altered behaviour. In this study, 73 non-depressed, high functioning, chronic, first-time stroke survivors performed a simple warned and unwarned auditory reaction time task. We show a reduced suppression of corticospinal excitability during movement preparation, an increased facilitation immediately prior to movement onset and slower reaction times in those with greater levels of PSF. Reduced suppression and increased facilitation of corticospinal excitability prior to movement onset in high fatigue is an indicator of poor modulation of pre-movement excitability which may in turn reflect poor sensory processing, supporting the sensory attenuation model of fatigue.

## Introduction

Post-stroke fatigue (PSF) is one of the most commonly self-reported symptoms after stroke that has significant implications for morbidity, disability, quality of life and mortality (Barbour and Mead, 2012). The majority of stroke survivors report fatigue in the first few weeks after stroke. This is linked to high levels of inflammation immediately following an injury and such fatigue is part of sickness behaviour (Dantzer and Kelley, 2007). The more debilitating symptom however, is fatigue that fails to resolve and persists for months or even years after the stroke. Despite the high prevalence of PSF, an understanding of the underlying mechanisms that underpin PSF is currently lacking (De Doncker et al., 2018).

We have previously shown that self-selected ballistic movement times are slower and resting corticospinal excitability assessed using transcranial magnetic stimulation (TMS) is reduced in the affected hemisphere of stroke survivors who report high levels of PSF (A. Kuppuswamy et al., 2015; Annapoorna Kuppuswamy et al., 2015). Speed of ballistic movements are dependent on the ability of the motor cortex to activate necessary corticospinal output to initiate movement and are intrinsically linked to corticospinal excitability (Jäncke et al., 2004). The state of the motor cortex at a time prior to movement, commonly referred to as the ‘movement preparation’ period plays a crucial role in determining ballistic movement speeds. Notably, corticospinal excitability does not only change during movement initiation but also undergoes distinct modulation during movement preparation (Cisek and Kalaska, 2010).

When preparing for a voluntary movement, there are substantial changes in the activity of neurons within the premotor and motor cortex (M1) despite no electromyographic (EMG) activity (Tanji and Evarts, 1976). Movement preparation has been extensively studied in humans using TMS over M1 to probe corticospinal excitability changes during cue-driven reaction time (RT) paradigms (Duque and Ivry, 2009; Hannah et al., 2018; Hasbroucq et al., 1997; Ibáñez et al., 2019). Cues for guiding movement are probabilistic in nature and learning the probabilities of upcoming movements enable the motor system to prepare motor output prior to movement initiation. Suppression of corticospinal excitability prior to movement initiation is seen in muscles that are both involved and uninvolved in an action (Bestmann and Duque, 2016; Duque et al., 2017; Greenhouse et al., 2015). The predicted aspects of sensory information (target cues in RT paradigms) are represented explicitly in the modulation of corticospinal excitability during movement preparation. One of the proposals of sensory attenuation model of fatigue is that, the inability to suppress predicted sensory stimuli results in high perceived effort leading to fatigue (Kuppuswamy, 2017). Therefore, pre-movement excitability representing predicted sensory information may be altered in those who exhibit high levels of fatigue. Behaviourally, such changes may result in reduced movement speeds (mediated by high perceived effort) which has been seen in stroke survivors with high fatigue (A. Kuppuswamy et al., 2015). Indeed, such differences in corticospinal excitability during movement preparation have previously been reported in multiple sclerosis patients with high and low fatigue (Morgante et al., 2011).

Therefore, the aim of this study was to investigate the modulation of corticospinal excitability leading up to a movement and reaction times in stroke survivors with varying severity of PSF.

## Results

### Demographics

There was a significant positive association between FSS-7 and anxiety (rho = 0.39, p < 0.01), depression (rho = 0.52, p < 0.01) and reaction time (rho = 0.25, p = 0.03), figure 2a-c. There was no association between FSS-7 and age nor RMT. FSS-7 was significantly greater for females (median = 4.57) than for males (median = 3.29), W = 418.5, p = 0.047 (figure 2d). There was no difference in FSS-7 in left hemisphere strokes compared to right hemisphere strokes.

**Figure 1.**
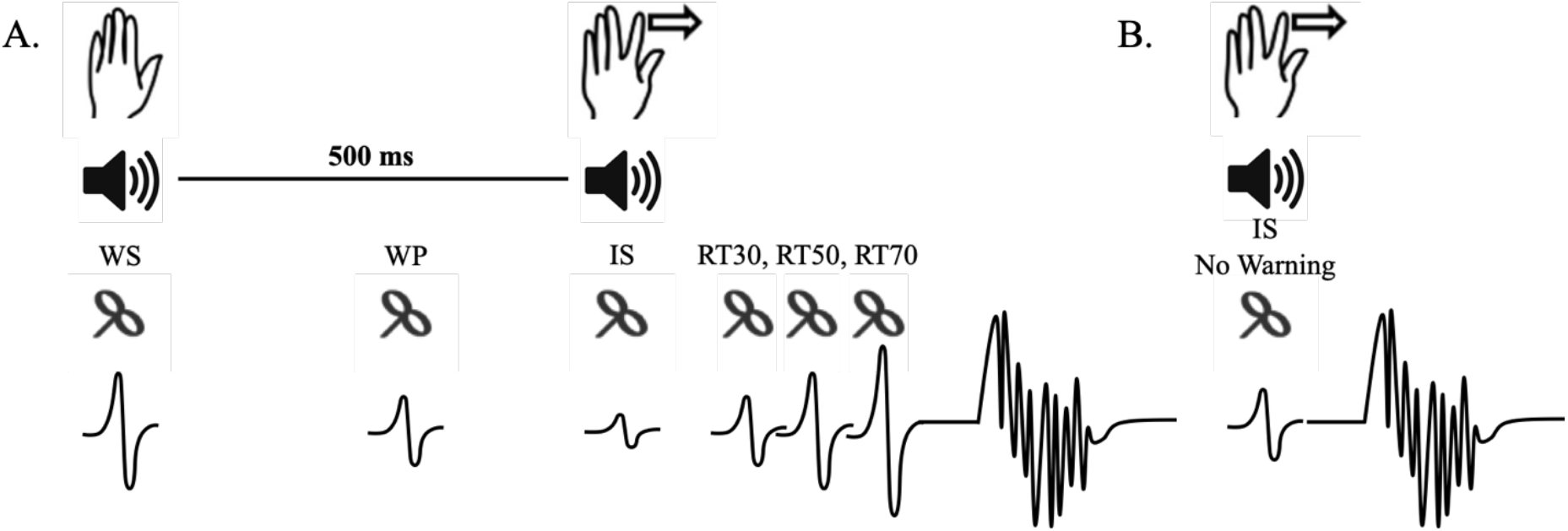
Study design for the simple auditory warned reaction time task (**A**) and the auditory unwarned reaction time task (**B**) used to study the modulation of corticospinal excitability during movement preparation. WS, WP, IS, IS No Warning, RT30, RT50 and RT70 indicate the different time points of stimulation across each task. Participants were instructed to carry out a ballistic index finger abduction after the auditory IS.

**Figure 2.**
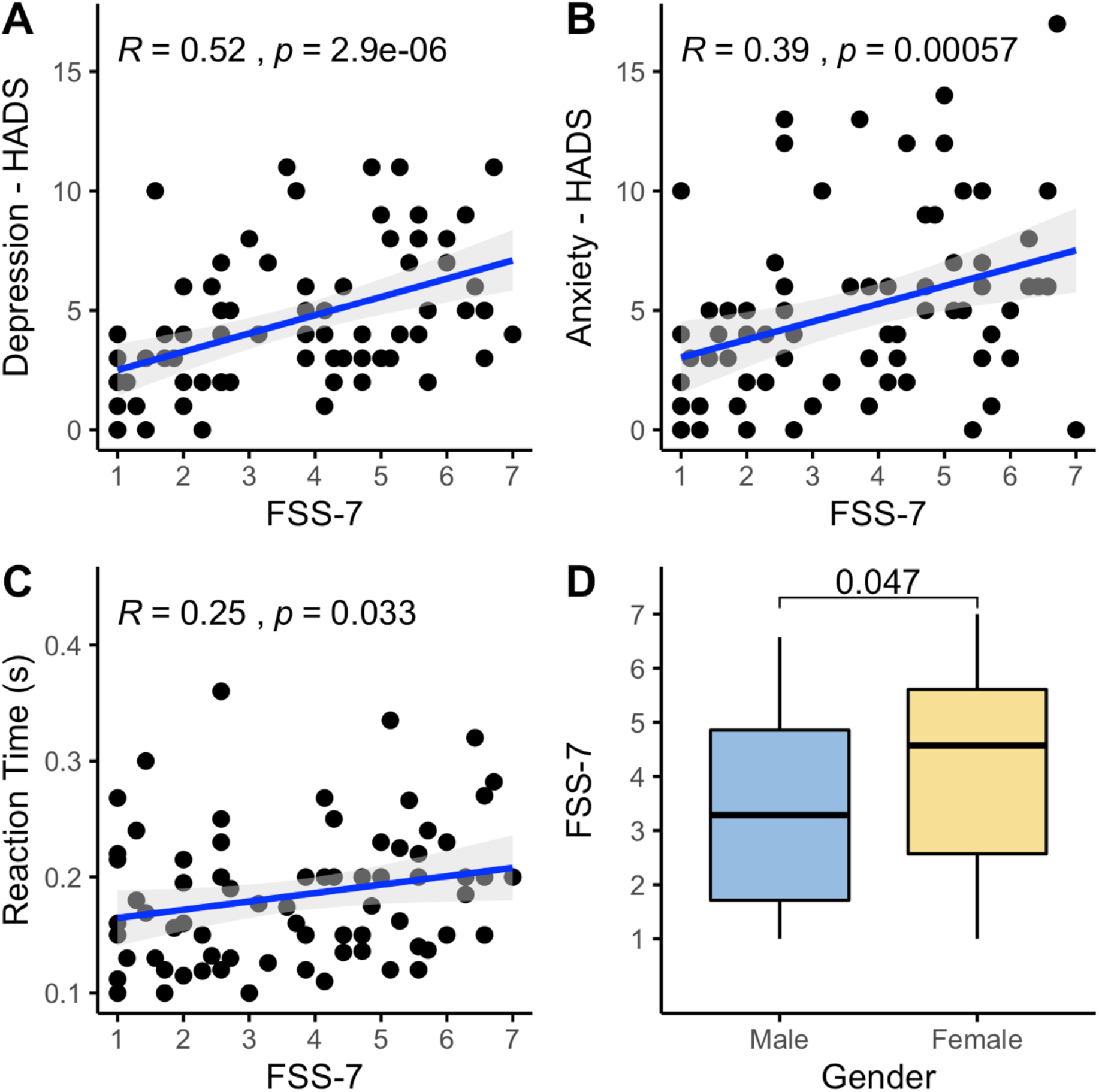
Associations between fatigue as measured by the FSS-7 and baseline measures such as depression (**A**), anxiety (**B**), baseline reaction time (**C**) and gender (**D**).

### Corticospinal excitability

A non-linear (quadratic) mixed effects model with time (WS, WP, IS, RT30, RT50 and RT70) and FSS-7 as fixed effects and participant nested in time as random effects best described the rate of change of corticospinal excitability during movement preparation. Including covariates that significantly correlated with FSS-7 (anxiety, depression, baseline reaction time and gender) did not significantly improve the model. The mixed effects model showed that time^2^ was a significant predictor of MEP amplitude (β = −0.44, t = −4.73, p < 0.001), FSS-7 did not significantly predict MEP amplitude (β = −0.068, t = 0.321, p = 0.75) and the interaction between time^2^ and FSS-7 was a significant predictor of MEP amplitude (β = −0.0066, t = −2.22, p = 0.0263) such that stroke survivors with higher fatigue showed less modulation of corticospinal excitability during movement preparation. There was a significant positive correlation between FSS-7 and MEP amplitude across all time points (WS: rho = 0.1, p = 0.02, WP: rho = 0.13, p = 0.003, IS: rho = 0.21, p < 0.001, RT30: rho = 0.18, p = 0.006, RT50: rho = 0.20, p = 0.003, RT70: rho = 0.13, p = 0.005). All corticospinal excitability data is presented in figure 3.

**Figure 3.**
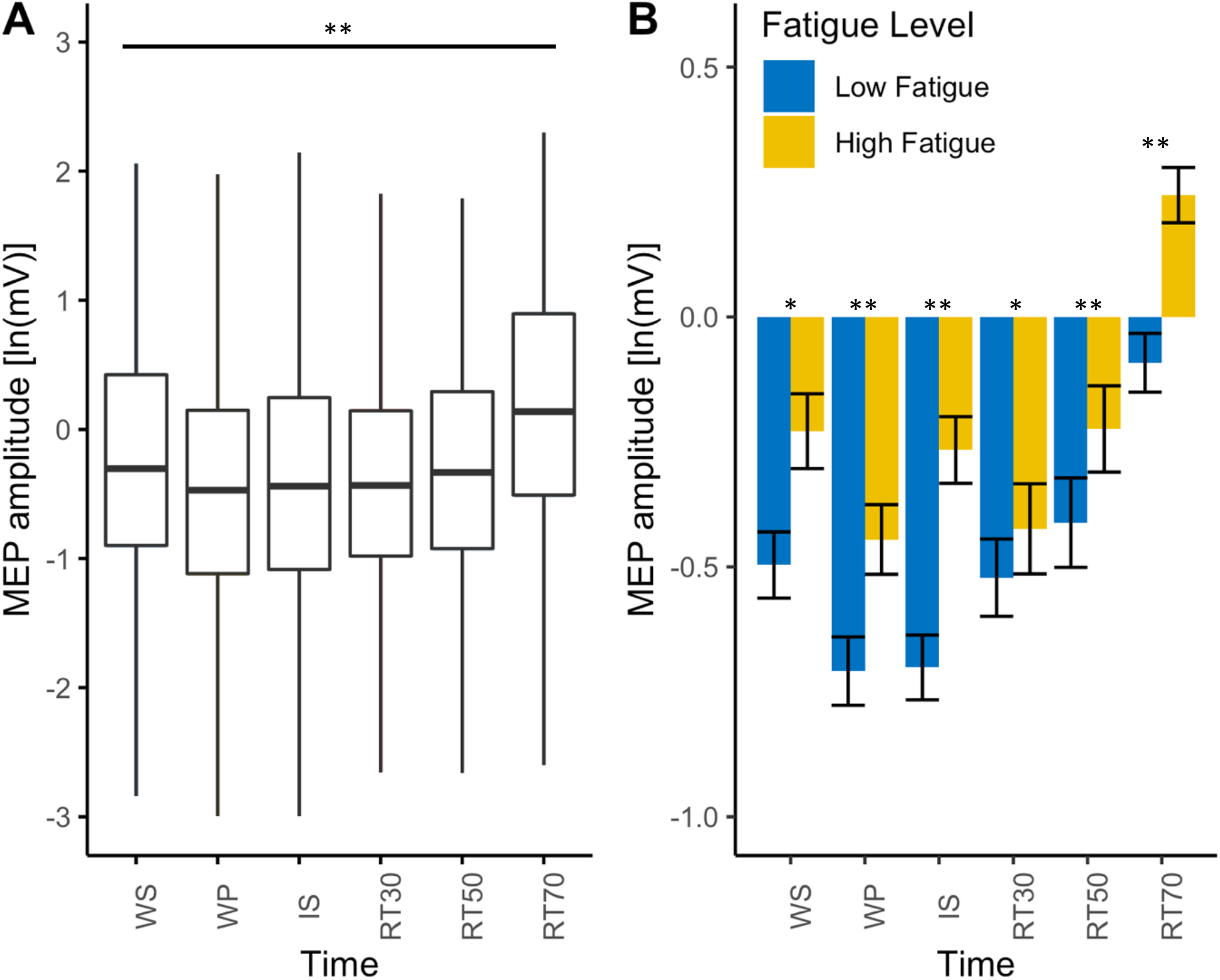
The effect of time and fatigue on MEP amplitude. **A.** boxplot for MEP amplitude across all stroke survivors for each time point indicating the significant effect of time on MEP amplitude. **B.** barplots with standard error bars representing MEP amplitude across all time points with stroke survivors grouped based on their fatigue score indicating the significant interaction between time and fatigue on MEP amplitude. Low fatigue patients (FSS-7 < 4) are represented in blue and high fatigue patients (FSS > 4) are represented in yellow. The association between fatigue (FSS-7) and MEP amplitude for each time point is also shown. Significance levels are indicated by * (* < 0.05, ** < 0.001).

### Reaction Time

A linear mixed effects model with time (WS, WP, IS, RT30, RT50 and RT70) and FSS-7 as fixed effects and participant nested in time as random effects best described the rate of change of reaction time during movement preparation. Including covariates that significantly correlated with FSS-7 (anxiety, depression, baseline reaction time and gender) did not significantly improve the model. The mixed effects model showed that the fixed effects of time (β = 0.0039, t = 0.980, p = 0.363) and FSS-7 (β = −0.0033, t = −0.854, p = 0.396) did not significantly predict RT. The interaction between time and FSS-7 (β = 0.0024, t = 2.47, p = 0.0159) was a significant predictor of RT such that stroke survivors with higher fatigue showed slower RTs the closer the stimulation time to movement onset. There was a significant positive correlation between FSS-7 and RT only at the IS (rho = 0.1, p = 0.01), RT50 (rho = 0.31, p < 0.001) and RT70 (rho = 0.32, p < 0.001) time points. All RT data is presented in figure 4.

**Figure 4.**
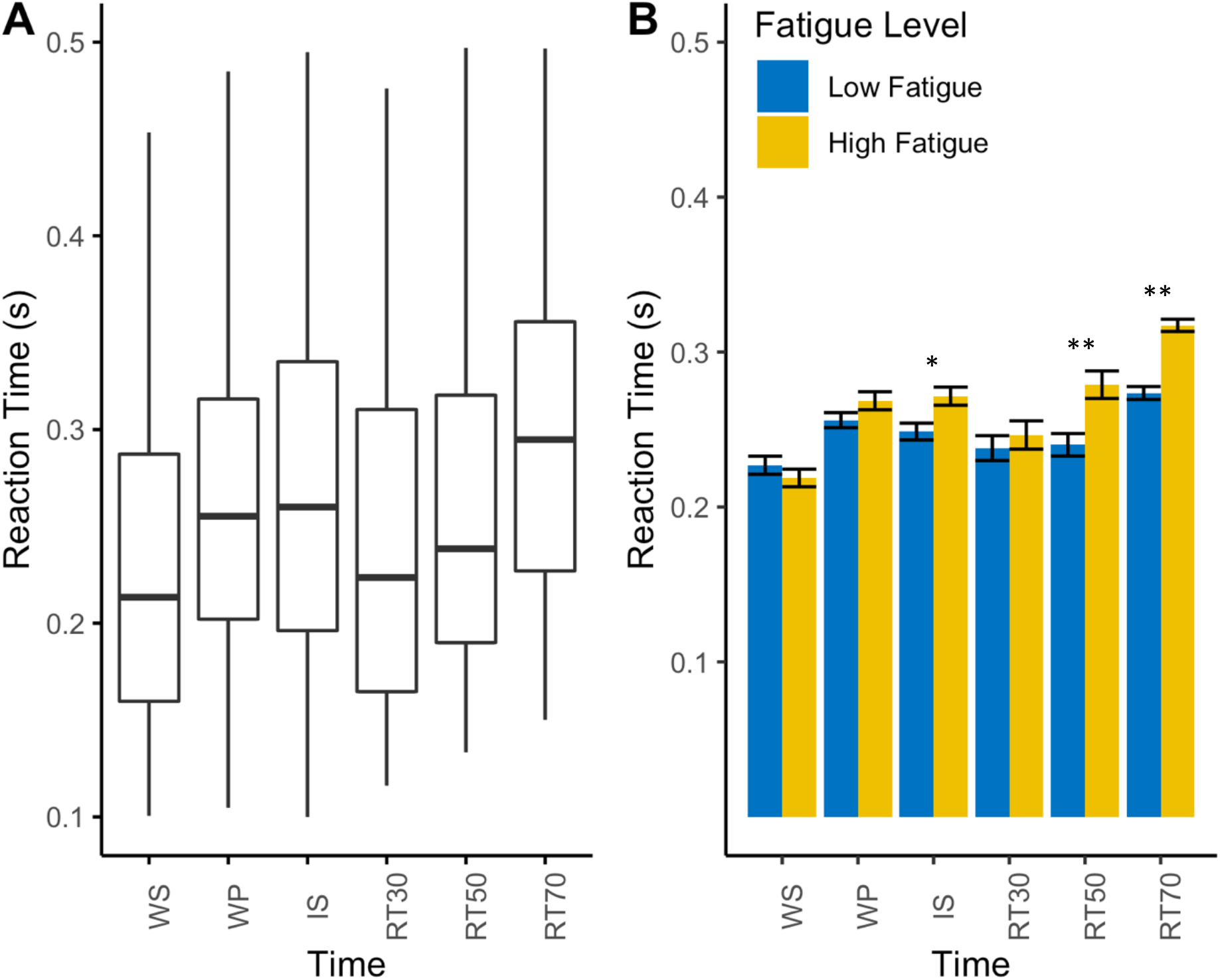
The effect of time and fatigue on RT. **A.** boxplot for RT across all stroke survivors for each time point indicating no significant effect of time on RT. **B.** barplots with standard error bars representing RT across all time points with stroke survivors grouped based on their fatigue score indicating the significant interaction between time and fatigue on RT. Low fatigue patients (FSS-7 < 4) are represented in blue and high fatigue patients (FSS > 4) are represented in yellow. The association between fatigue (FSS-7) and MEP amplitude for each time point is also shown. Significance levels are indicated by * (* < 0.05, ** < 0.001).

### Effect of Warning

A linear mixed effects model with condition (Warning, No Warning) as a fixed effect and participant nested in condition as random effects best described the difference in corticospinal excitability and RT across the two conditions. FSS-7 did not significantly improve the model in either of the two cases. The mixed effects model showed that the fixed effect of condition (β = 0.101, t = 4.072, p < 0.001) was a significant predictor of MEP amplitude across the two conditions (figure 5a). The fixed effect of condition (β = −0.004, t = −1.92, p = 0.0592) was not a significant predictor of RT across the two conditions (figure 5b). There was a significant positive correlation between FSS-7 and MEP amplitude across both conditions (Warning: rho = 0.21, p < 0.001, No Warning: rho = 0.11, p = 0.004) and a significant positive correlation between FSS-7 and RT in the Warning condition (rho = 0.1, p = 0.001), figure 5c-d.

**Figure 5.**
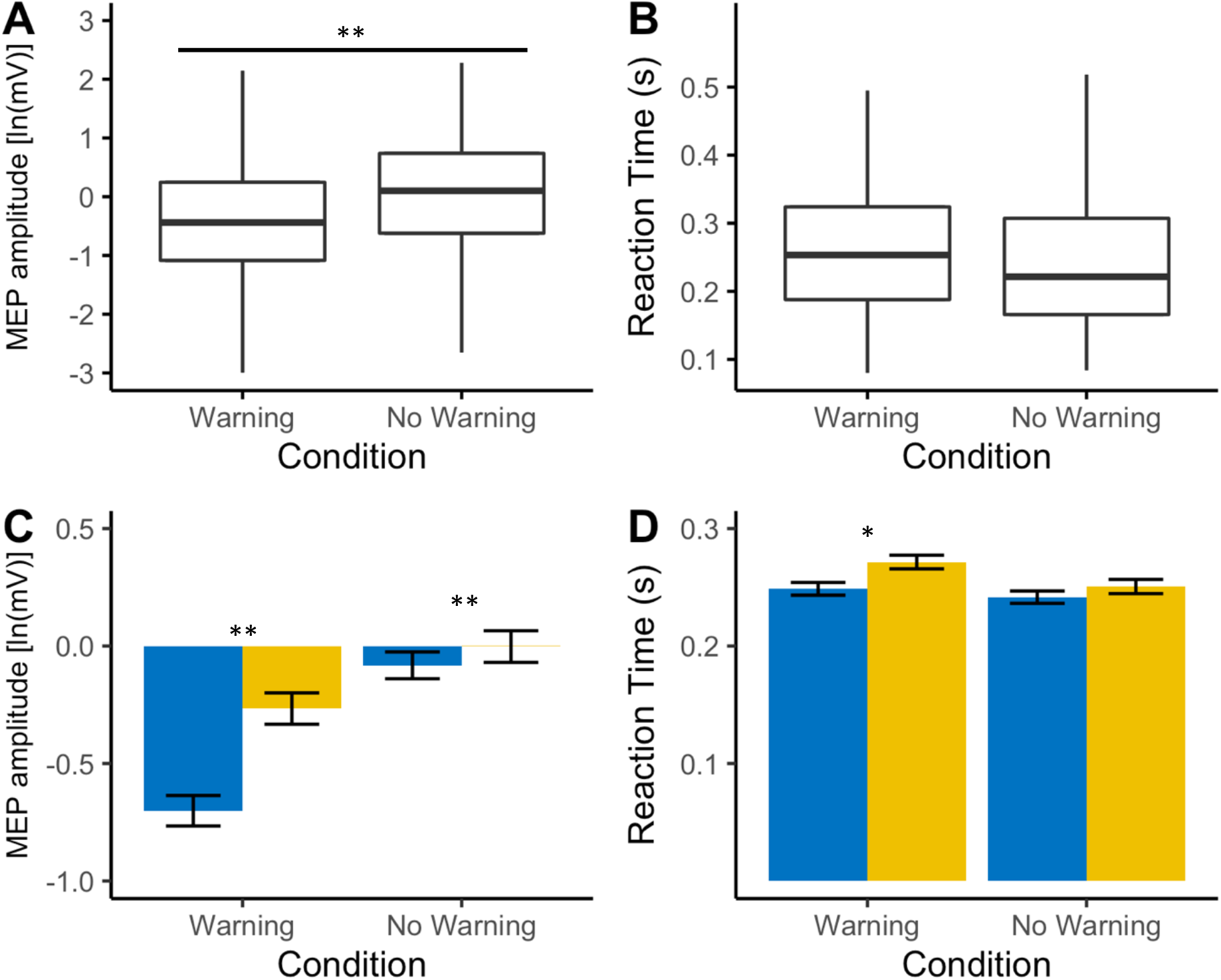
The effect of condition (Warning vs No Warning) and fatigue on MEP amplitude and RT. **A, B.** boxplots for MEP amplitude and RT across all stroke survivors for each condition indicating a significant effect of condition on MEP amplitude but not on RT. **C, D.** barplots with standard error bars representing MEP amplitude and RT across both conditions with stroke survivors grouped based on their fatigue score. Low fatigue patients (FSS-7 < 4) are represented in blue and high fatigue patients (FSS > 4) are represented in yellow. The association between fatigue (FSS-7) and MEP amplitude for each time point is also shown. Significance levels are indicated by * (* < 0.05, ** < 0.001).

## Discussion

In this study we showed that the modulation of corticospinal excitability during movement preparation changes as a function of fatigue in stroke survivors. Specifically, the higher the level of fatigue, the lower the suppression of corticospinal excitability during movement preparation and the higher the pre-movement facilitation of corticospinal excitability immediately before EMG onset. Reaction times were also greater in stroke survivors with greater fatigue during movement preparation. This was more apparent when TMS was delivered at time point closest to onset of EMG. We also showed that corticospinal excitability is higher in the absence of the warning cue in all stroke survivors. Analysis on the demographics of our patient cohort showed that females reported higher severity of fatigue when compared to males, and the higher the levels of anxiety and depression the higher the severity of fatigue. Stroke survivors with higher severity of fatigue also had slower reaction times at baseline.

A number of different models have been put forward to explain the reported modulation in corticospinal excitability during movement preparation from a motor control perspective (Burle et al., 2004; Duque et al., 2010; Duque and Ivry, 2009; Greenhouse et al., 2015; Lebon et al., 2019). Recent studies in nonhuman primates show that neural activity during movement preparation is a necessary component of movement generation and the time spent in the preparatory state can change depending on task demands (Lara et al., 2018). During cue driven movements, such as reaction time tasks where time is available, neuronal activity reaches the preparatory state well before movement onset. The ability to rapidly and consistently reach the correct preparatory state in advance, may allow time to correct small inaccuracies or even speed up movements before the movement itself (Churchland and Shenoy, 2007). Similar results have been reported in the mouse motor cortex as well as in humans, where the amount of preparatory inhibition, as measured by pre-movement cortical inhibition, positively correlated with reaction time (Hannah et al., 2018; Hasegawa et al., 2017). Stroke survivors with low levels of fatigue appear to follow a similar pattern of preparatory inhibition as healthy volunteers as has previously been described (Duque et al., 2017). However, with increasing fatigue, there is both reduced preparatory inhibition and earlier pre-movement facilitation before EMG onset. Reaction times are also slower with increasing severity of fatigue. Slower reaction times with greater fatigue, both at baseline and during the warned reaction time task maybe due to inability to reach the appropriate preparatory state, indicated by reduced preparatory inhibition, from which to initiate a movement. This could also explain why stroke survivors with high fatigue also have slower self-selected ballistic movement speeds (A. Kuppuswamy et al., 2015). Given the undue influence of the stimulation itself on reactions times, particularly at time points close to EMG onset, we were not able to examine the relationship between reaction times and corticospinal excitability in the current experiment. Whether the relationship between reaction times and changes in corticospinal excitability is causal should be explored further.

A number of studies have attempted to explain the changes reported in corticospinal excitability during movement preparation from a decision making and sensory processing perspective (Chiu et al., 2014; Cos et al., 2014; Freeman et al., 2014; Klein et al., 2012; Klein-Flügge et al., 2013; Klein-Flügge and Bestmann, 2012). MEP amplitudes may be influenced by other non-motor areas such frontal, parietal and subcortical regions that have projections to the premotor and motor cortex. Such influences include decision-related variables such as prior probabilities, subjective expected values or sensory evidence, which are computed elsewhere but ultimately influence the state of the motor system. When humans make choices between reaching actions, they tend to choose the one that is less effortful (Cos et al., 2014). This suggests that prior to movement initiation, one can predict the estimated action cost of different movements. Estimated action cost, normally experienced as ‘effort’, can therefore inform both implicit and explicit action choices towards the least effortful one. The estimated action cost of the upcoming movement is inversely proportional to the amplitude of MEPs (Cos et al., 2014). We have previously suggested that PSF is a result of altered perceptual processing, specifically altered perception of effort, associated with actions (Kuppuswamy, 2017). The results of the current study, altered modulation of corticospinal excitability, lend support to this hypothesis. The lack of preparatory inhibition and increased pre-movement facilitation may reflect a higher estimated action cost associated with the upcoming movement resulting in the movement being perceived as more effortful. Reduced pre-movement facilitation of corticospinal excitability in a visual reaction time task has also been reported in multiple sclerosis fatigue (Morgante et al., 2011). The authors conclude that impairment of areas engaged in motor planning might give rise to fatigue. This result provides further support to the hypothesis that fatigue is a result of altered effort perception, irrespective of disease pathology.

MEP amplitudes were lower in the warned RT condition compared to the unwarned RT condition irrespective of fatigue severity, while reaction times remained unchanged. Higher MEP amplitude in the unwarned condition is not surprising as the stimulation was given at the time of the imperative stimulus and movement “preparation” was yet to take place, unlike the warned condition. Given the difference in MEP amplitude between the two conditions we would also expect a difference in reaction times. One would expect reaction times to be slower in the unwarned condition when compared to the warned condition. The lack of difference in reaction times across the two conditions could be explained by the inclusion of catch trials in the warned condition experiment, whereas there were no catch trials in the unwarned condition. It has previously been reported that the presence of catch trials slows reaction times (Greenhouse et al., 2015). RTs appear to be longer in the warned condition with increasing severity of fatigue but unchanged in the unwarned condition. This might suggest that stroke survivors with high fatigue are capable of responding as quick as those with low fatigue when time is not available to prepare.

The current study further highlights the overlap of fatigue with other affective symptoms such as anxiety and depression previously described in the literature (De Doncker et al., 2018). Despite the overlap, the mechanisms underlying fatigue appear to be distinct as anxiety and depression scores did not improve the predictive capabilities of the model. This highlights the importance of using a strictly controlled patient cohort when trying to draw conclusions from studies and attempting to develop a mechanistic understanding of affective symptoms. In line with previous studies, we show that gender has an influence on incidence of fatigue, with females reporting higher levels of fatigue than males (Mead et al., 2011). However, this finding could be due to the fact that twice as many males than females took part in the study. Despite females reporting higher severity of fatigue as a group, gender did not appear to have any significant effect on corticospinal excitability and reaction times.

Despite providing us with useful insights into the mechanisms of post-stroke fatigue, this study is not without limitations. The intensity of stimulation to produce a 0.5mV response was determined at rest. However, when engaged in a task, corticospinal excitability at baseline, despite no EMG activity appears to be different across patients introducing variability into the paradigm. Future studies should ensure that all patients have a similar task-dependent baseline which they can use as a reference to compare the MEP amplitude across different time points. Given the nature of the symptom being investigated, a small number of trials was used at each time point to ensure that all patients could complete the task. We recommend that, given the variability of responses to TMS, future studies should use a block design with sufficient time between blocks to allow patients to rest in order to have a larger number of trials and more robust results. In the current study, we could not make a causal link between corticospinal excitability and reaction time due to the effect of the TMS pulse on reaction time. Using study designs in which reaction time is more closely controlled as demonstrated in a recent experiment (Ibáñez et al., 2019), we might be able to study the effect of pre-movement corticospinal excitability on reaction times.

To summarise, the modulation of corticospinal excitability during movement preparation assessed using TMS, changes as a function of fatigue in non-depressed, minimally impaired stroke survivors irrespective of anxiety and depression levels. Reaction times are also longer when given time to prepare for a movement in those who report high levels of fatigue. Preparatory inhibition, when viewed as a measure of sensory processing of expected stimuli, a reduction in preparatory inhibition in high fatigue may indicate poor sensory processing supporting the sensory attenuation model of fatigue.

## Materials and Methods

### Subjects

This is a cross-sectional observational study approved by the London Bromley Research Ethics Committee (REC reference number: 16/LO/0714). Stroke survivors were recruited via the Clinical Research Network from the University College NHS Trust Hospital, a departmental Stroke Database and from the community. All stroke survivors were screened prior to the study based on the following criteria: (1) first-time ischaemic or haemorrhagic stroke; (2) stroke occurred at least 3 months prior to the study; (3) no other neurological disorder; (4) not taking anti-depressants or any other centrally acting medication; (5) depression scores ≤ 11 assessed using the Hospital Anxiety and Depression Scale (HADS); (6) no sensory impairment; (7) grip strength and manual dexterity of the affected hand (at least 60% of the unaffected hand) assessed using a hand-held dynamometer and the nine hole peg test (NHPT) respectively; (8) no contraindications to TMS. Seventy-three stroke survivors took part in the study (demographics shown in table 1). All stroke survivors provided written informed consent in accordance with the Declaration of Helsinki.

**Table 1.** Demographics of all stroke survivors that took part in the study. Spearman’s rank correlation (Rho) and Wilcoxon rank sum tests (W) with their significance values for FSS-7 are reported for continuous and categorical data respectively.

|  | data (N = 73) | P-value |
| --- | --- | --- |
| **FSS-7** |  |  |
| Mean (SD) | 3.69 (1.84) |  |
| **Gender** | | $P = 0.0470$ |
| Females/Males | 24/49 |  |
| **Hemisphere Affected** | | $P = 0.3542$ |
| Left Hemisphere/Right Hemisphere | 40/33 |  |
| **Dominant Hand** | | $P = 0.0315$ |
| Right Hand/Left Hand | 67/6 |  |
| **Age (years)** |  |  |
| Mean (SD) | 61.55 (12.11) | $P = 0.1591$ |
| **Grip (% unaffected hand)** |  |  |
| Mean (SD) | 91.49 (18.82) | $P = 0.2836$ |
| **NHPT (% unaffected hand)** |  |  |
| Mean (SD) | 89.70 (23.66) | $P = 0.1497$ |
| **SDMT** |  |  |
| Mean (SD) | 1.19 (0.43) | $P = 0.5873$ |
| **Depression - HADS** |  |  |
| Mean (SD) | 4.56 (2.95) | $P < 0.0001$ |
| **Anxiety - HADS** |  |  |
| Mean (SD) | 5.04 (3.86) | $P = 0.0006$ |
| **RMT (% MSO)** |  |  |
| Mean (SD) | 47.97 (11.02) | $P = 0.4272$ |
| **Reaction Time (s)** |  |  |
| Mean (SD) | 0.18 (0.06) | $P = 0.0330$ |

### Surface Electromyogram and Transcranial Magnetic Stimulation

Recordings were carried out on the first dorsal interosseous (FDI) muscle of the affected hand. Following skin preparation using alcohol swabs, electromyogram (EMG) recordings were obtained from the FDI muscle using surface neonatal prewired disposable electrodes (1041PTS Neonatal Electrode, Kendell) in a belly-tendon montage with the ground positioned over the flexor retinaculum of the hand. The signal was amplified with a gain of 1000 (D360, Digitmer, Welwyn Garden City, UK), bandpass filtered (100-1000 Hz), digitized at 5kHz (Power1401, CED, Cambridge, UK) and recorded with Signal version 6.04 software (CED, Cambridge, UK). EMG recordings enabled the measurement of motor evoked potentials (MEPs).

A standard TMS device (Magstin 2002, Magstim, Whitland, Wales) connected to a figure-of-eight coil (wing diameter, 70mm) was used to stimulate the hand area of M1 in the hemisphere affected by the stroke. The coil was held tangentially on the scalp at an angle of 45° to the mid-sagittal plane to induce a posterior-anterior current across the central sulcus. The subjects were instructed to stay relaxed with their eyes open and their legs uncrossed. The motor ‘hotspot’ of the FDI muscle was determined as follows: the vertex (cross-over point between the mid-point between the two tragi and midpoint between nasion and inion) was marked using a dry wipe marker. Four centimetres lateral and 2 cm anterior from the vertex was then marked on the contralateral hemisphere, which is the approximate location of M1. This was used as a rough guide for a starting point for determining the hotspot for the first dorsal interosseous. At 50% MSO (maximal stimulator output) (or higher or lower in some patients) the coil was moved in 1 cm blocks for ~2 cm anterior, posterior, lateral and medial to the marked region. Three stimuli were delivered at each spot and the location with the highest average motor evoked potential response was taken as the hotspot.

### Resting motor threshold and 0.5mV intensity

Resting motor threshold (RMT) was defined as the lowest intensity of stimulation (% MSO) required to evoke a peak-to-peak MEP amplitude at the hotspot of at least 50μV in a minimum of 5 of 10 consecutive trials while subjects were at rest. Throughout the experiment, the stimulator setting was adjusted to produce a target MEP size of 0.5mV. This was defined as the stimulator setting (determined to the nearest 1% of MSO) required to evoke a peak-to-peak MEP amplitude of ≥ 0.5mV in a minimum of 5 of 10 consecutive trials.

### Simple Warned Reaction Time Task

Participants were seated comfortably in a chair facing away from the computer monitor with their hands palm-down on a pillow on their laps and performed a simple warned reaction time task. In each trial of the experiment, an auditory warning stimulus (WS) preceded an auditory imperative stimulus (IS) by a fixed interval of 500ms. Participants were instructed to respond quickly and accurately to the IS by making a finger abduction using the index finger of the hemiparetic side. Prior to the start of the experiment, participants completed 15 trials of the warned reaction time task without TMS in order to determine their mean baseline reaction time (RT). The main experiment consisted of a single block of 70 trials. To prevent anticipation of the IS and premature responses, catch trials were included where a WS was given with no IS and participants were instructed not to respond on these trials. TMS was delivered at five different time points (figure 1A): together with the WS (10 trials), late during the foreperiod defined as 167ms before the IS (10 trials), together with the IS (10 trials), at 30%, 50% and 70% of the mean baseline RT (10 trials for each) and at 70% of the mean baseline RT on the catch trials (10 trials). This resulted in six different conditions, which will be referred to as the following from now on: WS, WP, IS, RT30, RT50, RT70. This allowed us to measure corticospinal excitability during movement preparation. The order of trials was pseudorandomized across the five different TMS timings. Following the completion of 70 trials, a separate single block of 10 trials of unwarned reaction time task was completed. This was not included in the main experiment in order to maintain the effect of the WS. In this task, participants were given an auditory IS with no WS and were instructed to make a finger abduction using the same finger as previously described as quickly and accurately as possible. TMS was delivered together with the IS (figure 1B). In both experiments, stimulus timings were controlled using Signal version 6.04 software connected to a data acquisition system (Power1401, CED, Cambridge, UK). Each trial was 1.5 seconds long and the inter-trial interval was set to 5.5 ± 1.5 seconds.

### Fatigue

Fatigue was measured using the Fatigue Severity Scale (FSS-7), a widely used and validated questionnaire across different conditions ranging from 1-7 with an average score of seven being the highest fatigue and a score of one being no fatigue (Johansson et al., 2014).

### Data Processing and Statistical Analysis

#### Screening Test Scores

Spearman’s Rank Correlations between FSS-7 and a number of measures (age, anxiety, depression, grip strength, NHPT, SDMT, RMT and RT) were calculated. Wilcoxon rank sum tests were used to assess the difference in FSS-7 across different groups divided based on (gender, hemisphere affected and dominant hand being affected). The level of significance was set at p = 0.05.

#### TMS Data

The data files were extracted from Signal into MatLab and were analysed offline using custom-written routines in MatLab (2018a, Mathworks). Two dependent variables were measured on a trial-by-trial basis as follows: (1) MEP peak-to-peak amplitude and (2) reaction time (RT) measured from the time of the IS to the onset of volitional muscle activity. Peak-to-peak MEP amplitudes for each condition were estimated from the acquired EMG signal without applying any additional filters. Resting EMG was defined as the root mean square (rms) across all trials for each participant in the first 100ms of each trial (prior to the WS). Thresholds set at five times these levels were used to determine the RT. All trials were then visually inspected and manually corrected to ensure that RT was estimated properly and that no build-up of EMG was apparent before the TMS. Trials in which RT was less than 75ms or greater than 500ms were excluded from the final analysis as they represented premature and late responses respectively. Trials were also excluded if the MEP amplitude was less than 0.025mV. A logarithmic transformation (to the base of e) of MEP amplitudes was performed before the statistical tests to ensure normality of the samples. Trials containing outlier MEP amplitudes (Grubb’s test, p < 0.005) were also excluded from the final analysis.

To examine the effect of fatigue on corticospinal excitability and RT, log-transformed MEP amplitudes and RTs were labelled according to the time at which TMS was delivered (WS, WP, IS, RT30, RT50, RT70) and analysed by means of generalized mixed effects models carried out within the R environment for statistical computing, using the ‘lme4’ package (Bates et al., 2014). The ‘lmerTest’ package (Kuznetsova et al., 2017) was used to estimate the p-values for the t-test based on the Satterthwaite approximation for degrees of freedom. A stepwise ANOVA based on Satterthwaite’s approximation of degrees of freedom for model selection (lowest AIC value and p-value) was used to identify the combinations of variables that best predicted the outcome variables (MEP amplitude and RT). A similar analysis was used to examine the effect of fatigue and condition (Warning condition vs No Warning condition) on corticospinal excitability and RT. Assumptions of normality and homoscedasticity of the residuals for each model were assessed visually using quantile-quantile normal plots and fitted-versus residual-value plots. Individual spearman’s rank correlations were carried out between FSS-7 and the dependent variable in each model.

## Acknowledgments

We sincerely thank Prof John C. Rothwell and Dr Jaime Ibáñez for the many useful discussions and encouragement throughout the project. We thank Mr Cameron Cook and the Clinical Research Network for their help with recruitment. We thank our lab manager Mr Paul Hammond for the technical support throughout the project. We extend our heartfelt thanks to all our stroke survivor participants in this study without whose enthusiasm and commitment this study would not have been possible.

